# Daily-Academic: A Semantic Literature Discovery and Journal Tracking Platform for Biomedical Research

**DOI:** 10.64898/2026.11.19.767813

**Authors:** Dimitris Grigoriadis, Ilias Georgakopoulos-Soares, Georgios A. Pavlopoulos, Nikos Perdikopanis

## Abstract

**Objective:** Biomedical researchers monitoring newly published literature rely on fragmented combinations of keyword alerts, manual database searches, journal subscriptions and repository portals. We aimed to develop and evaluate an integrated platform that unifies semantic literature discovery, journal monitoring and linked omics dataset exploration within a single researcher-oriented workflow, driven by a user profile that remains explicit and editable.

**Methods:** Daily-Academic aggregates publication metadata from biomedical literature repositories and public omics resources and encodes titles and abstracts as dense semantic representations to support similarity-based retrieval beyond keyword matching. Recommendations are ranked by combining semantic relevance to the user profile with publication recency and journal-level preferences. The profile is derived from three user-supplied signals: a free-text statement of research interests, an optional link to the user’s own publication record, and interaction history. Retrieval quality was assessed on an offline benchmark of 30 biomedical queries over a fixed corpus of 259,246 publications, comparing Daily-Academic with a lexical BM25 baseline and a PubMedBERT bi-encoder under pooled assessment with graded relevance judgements.

**Results:** Query-level analysis indicated that the differences were distributed across queries rather than driven by a small subset.

**Conclusion:** Integrating semantic retrieval into a unified discovery workflow with an inspectable user model improved the quality of top-ranked biomedical literature relative to established lexical and dense baselines. Because the benchmark is user-free, the adaptive component of personalisation was inactive and the ranking weights untuned; the observed advantage therefore reflects the semantic representation rather than adaptive personalisation. The platform is publicly available at https://www.daily-academic.com.

## Introduction

Biomedical research output has grown at an unprecedented rate, with millions of new publications added annually across life science disciplines, placing increasing strain on researchers’ ability to monitor relevant findings in their fields. The growing volume of published scientific articles, datasets, and interdisciplinary research outputs has significantly increased the complexity of scientific information discovery and routine literature monitoring. Recent estimates suggest that millions of new scientific publications are added annually across biomedical and life science disciplines [1]. This continuous growth intensifies the operational and cognitive burden associated with identifying relevant and high-quality research findings.

Recent advances in biomedical informatics and AI-assisted scientific discovery have led to the development of diverse platforms supporting literature retrieval, semantic exploration, recommendation workflows, citation analysis, and evidence synthesis. Traditional biomedical literature discovery infrastructures, including PubMed [2] and Europe PMC [3], provide large-scale indexing and keyword-based retrieval of scientific publications. These resources form the foundation of modern biomedical literature access. Reference management and organizational platforms such as Zotero [4] and Mendeley [5] additionally support citation organization, journal monitoring, and personal literature management workflows commonly used by researchers.

More recent semantic and AI-assisted discovery systems have expanded beyond traditional keyword search toward personalized recommendation and relationship-based exploration of scientific content. Semantic Scholar [6] applies large-scale semantic analysis and machine learning techniques to support adaptive literature discovery and recommendation workflows. Platforms such as ResearchRabbit [7], Litmaps [8], and Connected Papers [9] focus on citation graph exploration and visual navigation of related scientific publications. Systems such as R Discovery [10] emphasize adaptive publication recommendation and personalized scientific monitoring workflows. Additional platforms including Scite [11] provide citation intelligence frameworks that classify and contextualize scientific citations to support literature interpretation and credibility assessment.

**Table 1.** Comparison of Daily-Academic with representative literature discovery platforms. The table summarizes the major functional characteristics of commonly used biomedical literature discovery systems and highlights the integrated workflow-oriented design of Daily-Academic. Feature comparison based on publicly available documentation and platform interfaces.

| Feature | Daily-Academic | R Discovery | ResearchRabbit | Semantic Scholar |
|---|---|---|---|---|
| Personalized recommendation feeds | ✓ | ✓ | Partial | ✓ |
| Semantic similarity-based retrieval | ✓ | ✓ | ✓ | ✓ |
| Journal and source tracking | ✓ | Limited | ✗ | Partial |
| Linked public omics dataset exploration | ✓ | ✗ | ✗ | ✗ |
| Interaction-aware personalization | ✓ | Partial | Partial | Limited |
| Integrated monitoring workflow support | ✓ | Partial | Partial | ✗ |
| Collaborative organization features | ✓ | Partial | Partial | ✗ |

Parallel advances in large language models and semantic reasoning have also led to the emergence of AI-assisted evidence synthesis and question-oriented scientific exploration systems. Platforms such as Elicit [12] and Consensus [13] leverage semantic retrieval and large language models to support evidence aggregation, scientific summarization, and question-answering workflows. In the biomedical domain, systems such as PubTator [14] incorporate biomedical entity recognition, semantic annotation, and literature enrichment techniques to facilitate structured exploration and knowledge extraction from large-scale biomedical publication corpora.

Despite these advances, many existing systems primarily focus on isolated aspects of scientific discovery, including semantic retrieval, citation exploration, evidence synthesis, reference management, or question answering. Fewer platforms emphasize integrated biomedical literature monitoring workflows that combine adaptive recommendation, structured journal tracking, semantic discovery, collaborative organization, and linked public omics dataset exploration within a unified environment optimized for rapid daily literature triage and longitudinal scientific monitoring.

Biomedical researchers therefore frequently continue to rely on fragmented combinations of keyword alerts, manual PubMed searches, journal subscriptions, reference management tools, and external dataset repositories to remain informed about newly published scientific findings and associated biological datasets. This fragmentation increases the operational and cognitive burden associated with routine literature monitoring. It may also reduce the efficiency of identifying relevant publications across rapidly evolving biomedical domains.

To address these challenges, we developed Daily-Academic, a biomedical informatics platform designed to support semantic literature discovery, adaptive scientific monitoring, and integrated biomedical research workflows.

Daily-Academic integrates publication metadata from biomedical literature repositories and public omics resources to support unified retrieval across scientific publications and associated biological datasets. Recommendations adapt over time as the platform incorporates each user’s reading behavior, saved articles, and journal preferences into an evolving interest profile.

These mechanisms facilitate rapid literature triage and monitoring of newly published scientific content. The platform additionally supports topic and tag-based exploration, collaborative sharing workflows, and integrated monitoring of selected biomedical journals and publication sources.

The main contributions of this work are summarized as follows:

1. Development of a semantic biomedical literature discovery platform integrating personalized recommendation and journal tracking workflows within a unified researcher-oriented environment.
2. A vector-based semantic retrieval framework combining publication metadata, adaptive personalization signals, and interaction-aware ranking mechanisms.
3. Integration of biomedical literature monitoring with linked public omics dataset exploration to support unified discovery across publications and associated biological resources.
4. An offline retrieval benchmark evaluating semantic retrieval quality against established lexical and dense information-retrieval baselines across 30 biomedical queries.

The remainder of this paper describes the system architecture and evaluation framework in detail, followed by evaluation results and a discussion of platform limitations and future directions.

### Statement of Significance

#### Problem or Issue

Biomedical researchers monitoring new literature must move between separate tools, keyword alerts, PubMed searches, journal subscriptions, reference managers and dataset repositories. This fragmentation costs time daily and causes relevant work to be missed.

#### What is Already Known

Semantic retrieval based on biomedical language models improves relevance over lexical search, and several discovery platforms offer recommendation, citation exploration or evidence synthesis. These functions remain distributed across systems, and the user models driving relevance are typically opaque and not directly editable by the researcher.

#### What This Paper Adds

We present Daily-Academic, which unifies personalized recommendation, topic feeds, journal tracking and linked omics dataset discovery in one workflow, driven by an explicit user profile the researcher can inspect and revise. We evaluate retrieval quality against lexical and dense baselines on 30 biomedical queries under pooled graded relevance assessment.

#### Who Would Benefit

Biomedical and translational researchers who monitor literature routinely, and developers of discovery and recommendation systems for scientific information.

## Methods

### System overview

Daily-Academic was developed to address the growing fragmentation of biomedical literature monitoring workflows. The platform aggregates newly published scientific literature and associated omics dataset metadata into a unified semantic discovery environment, with a primary focus on recently published content relevant to life sciences and bioinformatics research.

The system is organized around a researcher-centered workflow: incoming publications are first acquired and normalized from biomedical repositories, then transformed into dense semantic representations that enable similarity-based retrieval. These representations feed into a personalization layer that continuously refines each user’s interest profile based on interaction signals, journal preferences, and saved content. The resulting ranked feeds are delivered through a web-based interface designed for rapid daily literature triage.

This workflow is implemented across five architectural components: data acquisition and normalization, semantic representation learning, vector-based retrieval and ranking, adaptive user personalization, and user-facing feed generation (Figure 1), each described in detail in the subsections that follow.

**Table 2.** Main features of Daily-Academic. Summary of the core functionalities provided by the platform, including personalized recommendations, semantic search, filtering, article tracking, and user interaction support.

| Feature Category | Functionality | Description |
|---|---|---|
| Personalized Recommendations | AI-based article ranking | Recommends scientific articles based on user interests, semantic relevance, interaction history, and recency signals. |
| Semantic Search | Natural language query understanding | Supports semantic retrieval of publications using contextual understanding rather than simple keyword matching. |
| Literature Feed | Personalized academic stream | Provides continuously updated article recommendations tailored to each user profile. |
| Filtering System | Advanced filtering options | Allows filtering by research field, journal, publication date, keywords, and other metadata. |
| Article Tracking | Saved and monitored articles | Enables users to bookmark, revisit, and organize publications of interest. |
| User Interaction Modeling | Interaction-aware ranking | Incorporates user clicks, reading behavior, and preferences into recommendation scoring. |
| Recency Awareness | Time-sensitive prioritization | Promotes recently published and emerging scientific literature when relevant. |
| Journal-Based Scoring | Source quality integration | Considers journal-related information as part of the ranking process. |
| Search Interface | Interactive exploration | Provides a user-friendly interface for browsing, searching, and exploring scientific content. |
| Metadata Visualization | Structured publication information | Displays article metadata including title, authors, journal, abstract, and publication date. |
| Multi-domain Support | Broad scientific coverage | Supports literature discovery across multiple biomedical and scientific domains. |
| Web-based Accessibility | SaaS platform availability | Accessible through a web browser without requiring local installation. |
| Dynamic Recommendations | Adaptive recommendation updates | Continuously refines recommendations based on evolving user interactions and interests. |
| Query Assistance | Flexible search capabilities | Supports both keyword-based and semantic-style user queries. |
| Research Discovery | Identification of relevant studies | Helps researchers identify potentially relevant publications more efficiently than traditional manual search workflows. |

**Figure 1.**
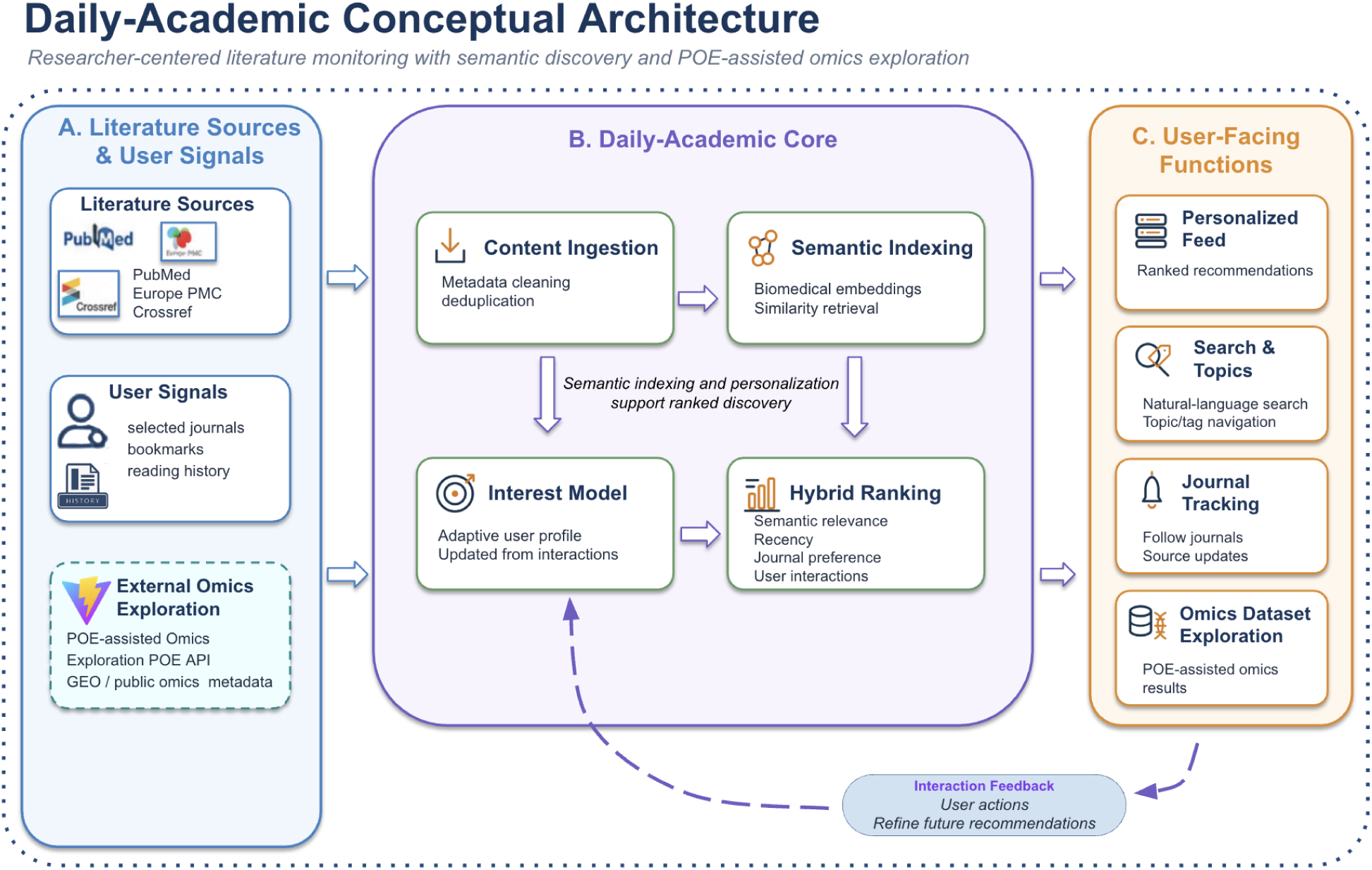
Conceptual architecture of Daily-Academic. The platform combines literature ingestion, semantic indexing, adaptive user-interest modeling, and hybrid ranking to support personalized biomedical literature monitoring. External literature sources and user interaction signals are processed within the Daily-Academic core, while POE-assisted omics exploration links publications to relevant public omics datasets. User-facing functions include personalized feeds, search and topic navigation, journal tracking, and omics dataset exploration. Interaction feedback is used to continuously refine future recommendations.

### Data Acquisition and Normalization

#### Scientific Literature Sources

Daily-Academic continuously aggregates newly published scientific literature from biomedical publication repositories, primarily PubMed. The ingestion pipeline is initiated by querying the Crossref API on a rolling 7-day window, filtering results against a curated set of approximately 3,000 manually selected journal ISSNs to ensure domain relevance and publication quality. The resulting DOIs are persisted in the database along with a datestamp for provenance tracking. A subsequent asynchronous routine resolves each DOI to its corresponding PubMed identifier (PMID) via the NCBI E-utilities (EFetch) API. Each PMID is then used to retrieve full bibliographic metadata from PubMed, including article title, authors, abstract, journal name, publication date, keywords, MeSH terms, figures, and publication type. To enrich article representations with visual content, the pipeline attempts to obtain a graphical abstract from ScienceDirect via its API, restricted to open-access content. In cases where no graphical abstract is available, the pipeline falls back to extracting the first figure by fetching the article’s DOI landing page. This step is silently skipped when the target page is inaccessible (e.g., rendered server-side via frameworks such as Firebase), ensuring pipeline robustness without interrupting the overall ingestion flow.

#### Omics Dataset Integration

In addition to scientific literature, the platform incorporates omics-related metadata through the Public Omics Explorer (POE) API [15]. POE aggregates and harmonizes metadata from public omics repositories, including the Gene Expression Omnibus, and exposes them to the platform as structured searchable records. These metadata include experiment type, such as RNA-seq or ATAC-seq, organism, phenotype or disease annotations, and accession identifiers, including GSE and SRR records.

A key component of the integration pipeline involves metadata enrichment and linkage between scientific publications and associated omics datasets. This linkage is established using publication identifiers, abstracts, MeSH annotations, and repository metadata. The resulting integration supports unified retrieval across biomedical literature and associated public datasets.

#### Content normalization

Following ingestion, publication and dataset records are transformed into a normalized internal representation. Normalization includes metadata harmonization, identifier cleaning, source labeling, and duplicate detection. Deduplication is particularly important because identical publications may appear through multiple indexing or hosting sources. Normalized records are subsequently stored for semantic indexing, ranking, filtering, and feed generation.

### Semantic Representation and Personalization

#### Semantic Embedding Generation

Scientific publications are encoded as 768-dimensional dense vector representations using a pretrained BioBERT-based SentenceTransformer model corresponding to the Hugging Face checkpoint pritamdeka/BioBERT-mnli-snli-scinli-scitail-mednli-stsb. The model has been fine-tuned for sentence-level semantic similarity and natural language inference tasks across general, scientific, and biomedical domains [16,17]. These embeddings support semantic similarity search by representing biomedical texts in a dense vector space that captures contextual and domain-specific relationships.

Prior to embedding, publication records are transformed into metadata-enriched textual representations. Long texts are handled through truncation or segment-level embedding followed by vector averaging, producing consistent document-level semantic representations within the transformer input-length constraints.

The platform does not further fine-tune the embedding model. Instead, embeddings are generated and indexed continuously as newly published literature and datasets are incorporated into the platform.

#### User Interest Representation

Personalization in Daily-Academic is based on signals reflecting user-specific scientific interests. These signals include selected journals, saved publications, dismissed items, interaction history, and platform activity patterns.

Users may optionally provide publication history information to initialize a preliminary representation of research interests.

Each user is represented through a continuously updated user-interest embedding vector. User representations are refined dynamically as the user interacts with recommended content. Incoming publications are compared against these adaptive representations to prioritize content within personalized recommendation feeds.

### Semantic Retrieval and Ranking

#### Similarity Retrieval

Semantic retrieval is performed using vector similarity search over dense document embeddings. Document relevance is estimated using cosine similarity:

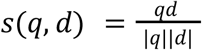

where q represents the user-interest embedding and d represents the document embedding.

This semantic retrieval framework enables identification of conceptually related biomedical publications and datasets even when exact terminology differs across documents.

Approximate nearest-neighbor retrieval is implemented using the FAISS similarity search framework [18]. Dense embedding vectors and associated relational metadata are stored within a PostgreSQL [19] database enhanced with the pgvector extension [20], enabling efficient vector-based retrieval alongside structured metadata management. This architecture supports unified document discovery through combined semantic similarity retrieval and metadata-aware filtering operations.

#### Ranking Framework

For each user, Daily-Academic generates personalized literature feeds using a hybrid ranking framework integrating semantic relevance, journal weighting, publication recency, and interaction-derived personalization signals.

Candidate publications are ranked using a weighted hybrid scoring function combining semantic relevance, source preferences, temporal prioritization, and interaction-derived personalization signals.

The overall ranking score R for a candidate document is computed as:

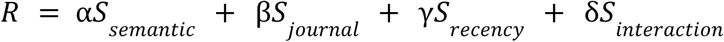

where:

- S_semantic_ represents the semantic similarity between the user-interest embedding and the document embedding;
- S_journal_ represents journal or source preference weighting derived from user-selected or frequently accessed publication venues;
- S_recency_ represents a time-sensitive recency component prioritizing recently published scientific literature;
- S_interaction_ represents interaction-derived personalization signals based on user activity patterns, including saved articles, clicks, and positive engagement behavior.

Individual score components are normalized prior to weighted aggregation to maintain comparable contribution scales across ranking dimensions.

The weighting parameters α, β, γ, and δ determine the relative contribution of each component and are empirically adjusted to balance semantic relevance, personalization, and recency-aware literature monitoring behavior. Additional filtering stages exclude duplicate records, previously reviewed publications, and content outside user-defined filtering criteria. The ranking framework is designed to support efficient daily scientific literature monitoring and rapid relevance-oriented content triage rather than exhaustive document retrieval.

### Infrastructure and Platform Implementation

#### Backend Infrastructure

The backend infrastructure of Daily-Academic follows a modular microservice-oriented architecture designed to support scalable biomedical literature ingestion, semantic retrieval, and personalized feed generation workflows. The platform integrates semantic vector representations with structured relational metadata to enable unified retrieval across biomedical publications and associated public omics datasets.

Publication and dataset embeddings are stored alongside relational metadata within a PostgreSQL database augmented with the pgvector extension, enabling simultaneous querying over structured metadata fields and semantic similarity representations. Database access is mediated through an object-relational mapping layer with connection pooling, ensuring transactional consistency and efficient concurrent access across ingestion and retrieval workloads. Semantic retrieval and approximate nearest-neighbor search are implemented using the FAISS similarity search framework, which supports efficient retrieval over high-dimensional embedding spaces.

The API layer is implemented using FastAPI [21] and supports asynchronous request handling and scalable feed generation workflows. Cross-origin resource sharing and response compression middleware are applied at the application entry point to ensure secure, low-overhead client–server communication. Newly published biomedical literature and associated omics metadata are continuously acquired through scheduled crawling and ingestion pipelines. Semantic indices are updated regularly to incorporate newly ingested content and maintain low-latency semantic retrieval.

Asynchronous background processing pipelines are used for crawling, embedding generation, feed pre-computation, and indexing tasks. Additional caching and asynchronous task scheduling mechanisms are employed to improve retrieval efficiency and reduce redundant feed generation operations for users with stable preference representations.

The platform includes an authentication and user-management subsystem that supports secure user access and personalized functionality. User identity is handled through Clerk [22], enabling federated sign-in via external providers such as Google, LinkedIn, and GitHub. Authenticated user profiles and personalization preferences are stored in relational database tables, allowing the platform to maintain user-specific recommendations, access control, and privacy-aware management of interaction data. To reduce redundant computation during repeated user sessions, a Redis-based caching layer [23] stores precomputed recommendation and similarity results, which are refreshed when new content is ingested or when user interaction patterns change.

#### User Interface and Workflow Design

The Daily-Academic user interface is implemented as a web-based single-page application using React [24], TypeScript [25] and Tailwind CSS [26]. The interface is designed to support low-friction biomedical literature monitoring, rapid relevance-oriented scientific content triage, and efficient daily review workflows across both desktop and mobile environments.

Users interact with organized recommendation feeds composed of article cards containing structured metadata designed to facilitate rapid relevance assessment and decision-making. Displayed metadata include article title, abstract snippet, publishing journal, publication date, article type (e.g., Review, Clinical Trial), open-access status, journal impact factor, and geo-location information associated with author affiliations. Each result additionally includes direct links to the original publication source and associated external biomedical database records.

Personalized recommendation relevance is visually represented through interface indicators highlighting publications with high semantic similarity to the user-interest profile representation. The platform additionally supports topic and tag-based exploration, collaborative article sharing workflows, and structured monitoring of selected biomedical journals and publication sources.

To support integrated scientific workflows, Daily-Academic includes Zotero integration, social sharing functionality, and collaborative user Groups that enable researchers to organize and exchange relevant scientific publications within defined cohorts. Journal tracking feeds are integrated directly into the broader recommendation environment, allowing users to combine personalized semantic discovery with source-specific literature monitoring within a unified interface.

The overall workflow is designed to support rapid review, filtering, saving, dismissal, and revisiting of scientific publications while minimizing the operational burden associated with fragmented biomedical literature monitoring workflows.

### Ethics, Privacy, and Data Handling

Daily-Academic was designed with attention to privacy-aware handling of user interaction and personalization data. The platform stores only the information required to support semantic recommendation, journal tracking, and personalized literature monitoring workflows.

User interaction signals, including saved articles, clicks, reading activity, and journal preferences, are used exclusively for adaptive recommendation and personalization within the platform. Personally identifiable information is minimized and is not shared with external third parties for advertising or commercial profiling purposes.

The platform primarily operates on publicly available scientific publication metadata and publicly accessible omics dataset metadata obtained from external repositories and APIs. No protected patient-level biomedical data or controlled-access genomic datasets are stored or processed within the current platform implementation.

Authentication and user-specific personalization workflows are implemented using secure access-control mechanisms. User activity data are processed within isolated user contexts to support individualized recommendation generation while reducing unnecessary exposure of interaction histories.

## Results

### Platform Overview

Daily-Academic provides an integrated biomedical literature discovery environment accessible through a web-based interface. The platform is designed to support rapid daily literature review and reduce the fragmentation associated with conventional biomedical literature monitoring workflows. The system combines personalized recommendation feeds, semantic retrieval, structured journal monitoring, topic exploration, and linked public omics dataset discovery within a unified environment tailored for biomedical and life science research.

### Personalized Recommendation Feed

The primary workflow is centered around a personalized recommendation feed that continuously surfaces newly published scientific literature ranked according to semantic relevance, publication recency, journal preferences, and interaction-derived personalization signals (Figure 2). Publications are presented through structured article cards displaying:

- Article title and abstract snippet
- Publishing journal and publication date
- Article type (e.g., Research Article, Review, Clinical Trial)
- Journal impact factor and open-access status
- Direct links to the original publication and associated external biomedical resources

**Figure 2.**
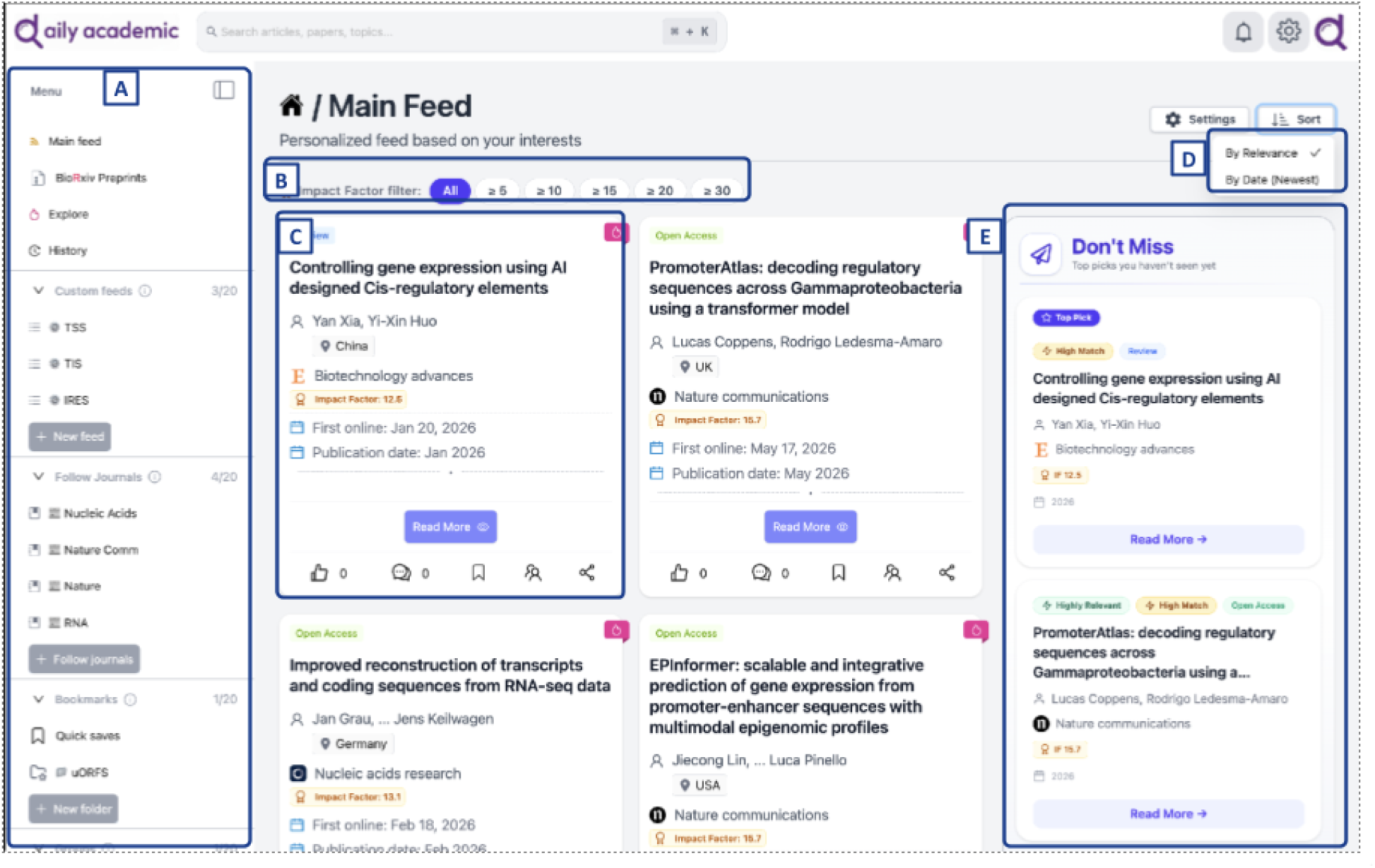
Personalized recommendation interface of Daily-Academic. (A) Navigation sidebar providing access to the main feed, preprint stream, user-defined custom feeds, followed journals, bookmark folders and collaborative groups. (B) Impact-factor threshold filter for on-the-fly restriction of the feed. (C) Article cards presenting title, authors, corresponding-author country, journal, impact factor and first-online/publication dates, together with viewing state and interaction controls (like, comment, bookmark, share). (D) Ranking control allowing the feed to be ordered either by semantic relevance to the user profile or by recency. (E) The “Don’t Miss” panel, which surfaces high-scoring articles the user has not yet opened, each annotated with a semantic match indicator.

Semantic relevance to the user’s interest profile is visually indicated for each article, enabling rapid relevance assessment without opening individual publications. Interaction controls allow users to save, dismiss, or rate articles, with each interaction refining subsequent recommendation output.

An advanced filtering panel supports narrowing of recommendation feeds by time range, publication type, journal or source, keywords, and author name. Feed tabs additionally allow switching between all recommendations, unread items, topic-specific views, and top journal outputs within a single interface.

### User-Controlled Personalization

Personalization is driven by three user-supplied signals that remain visible and editable at all times (Figure 3). Users describe their research interests in free text, from which the profile embedding is derived; they may optionally link an external author profile to initialize those interests from their own publication record; and they may enable an adaptive component that additionally updates the profile vector from articles they have upvoted (Supplementary Figure S1). This design keeps the basis of every recommendation inspectable and revisable, in contrast to systems in which the user model is implicit.

**Figure 3.**
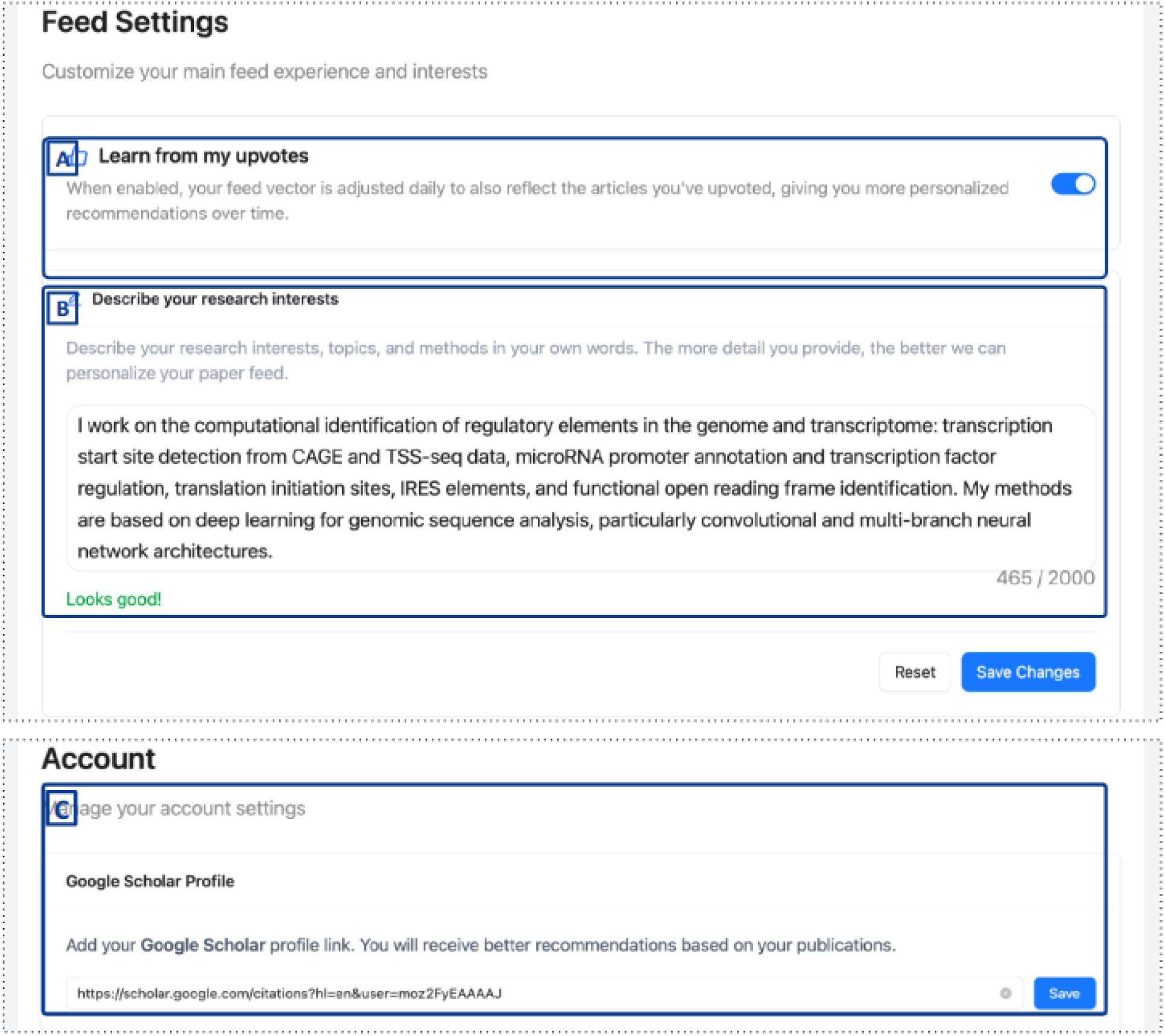
User-controlled personalization settings. (A) Optional adaptive component that updates the user profile vector from articles the user has upvoted. (B) Free-text description of research interests, from which the profile embedding is derived. (C) Linking of an external author profile, used to initialize interests from the user’s own publication record. Together these provide explicit, inspectable and revisable control over the signals driving recommendation.

### Custom Topic Feeds

Beyond the main feed, users can define persistent topic-specific feeds that are continuously updated as new literature is indexed (Figure 4). Each custom feed maintains its own article count and supports the same impact-factor filtering and card layout as the main view, allowing focused surveillance of a narrow research topic alongside general personalized discovery.

**Figure 4.**
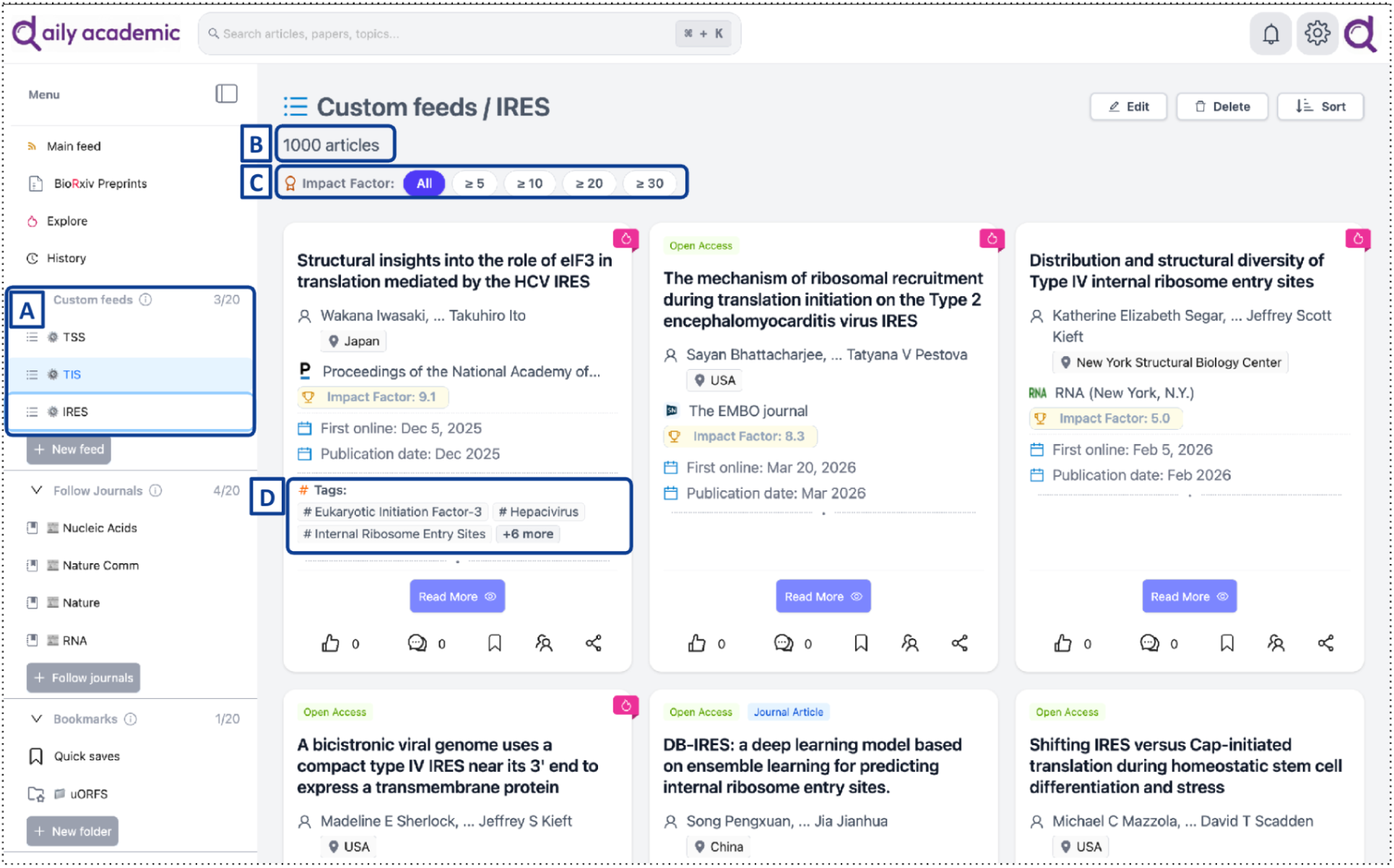
Topic-specific monitoring through user-defined custom feeds. (A) Custom feeds are listed alongside the main feed in the navigation sidebar. (B) Each feed reports the number of matching articles and is maintained as a persistent, continuously updated stream. (C) Impact-factor filtering operates within the feed as in the main view. (D) Indexed subject terms are shown on each card. The feed illustrated here tracks internal ribosome entry sites.

### Semantic Search and Topic Exploration

The platform supports semantic search across the indexed publication corpus using natural language queries, enabling retrieval of conceptually related publications beyond exact keyword matching. Search results can be filtered by article type, journal, author, and publication date, and are sortable by relevance or recency.

Topic and tag-based exploration allows users to navigate across semantically related biomedical research areas and emerging publication clusters. Topics are organized hierarchically into broad research areas, domain-specific subtopics, methodological categories, application areas, and dataset types. Each topic view displays associated article counts, trend data, related topics, and popular tags. User interactions with topics and tags are incorporated into the personalization model, progressively refining recommendation relevance as exploration activity accumulates.

### Journal Tracking

The journal tracking workflow enables users to follow selected biomedical journals and publication sources directly within the platform (Figure 5). Users can search and browse journals by name, subject area, or impact factor, and configure per-journal tracking preferences including notification frequency, topic filtering, and article type selection.

**Figure 5.**
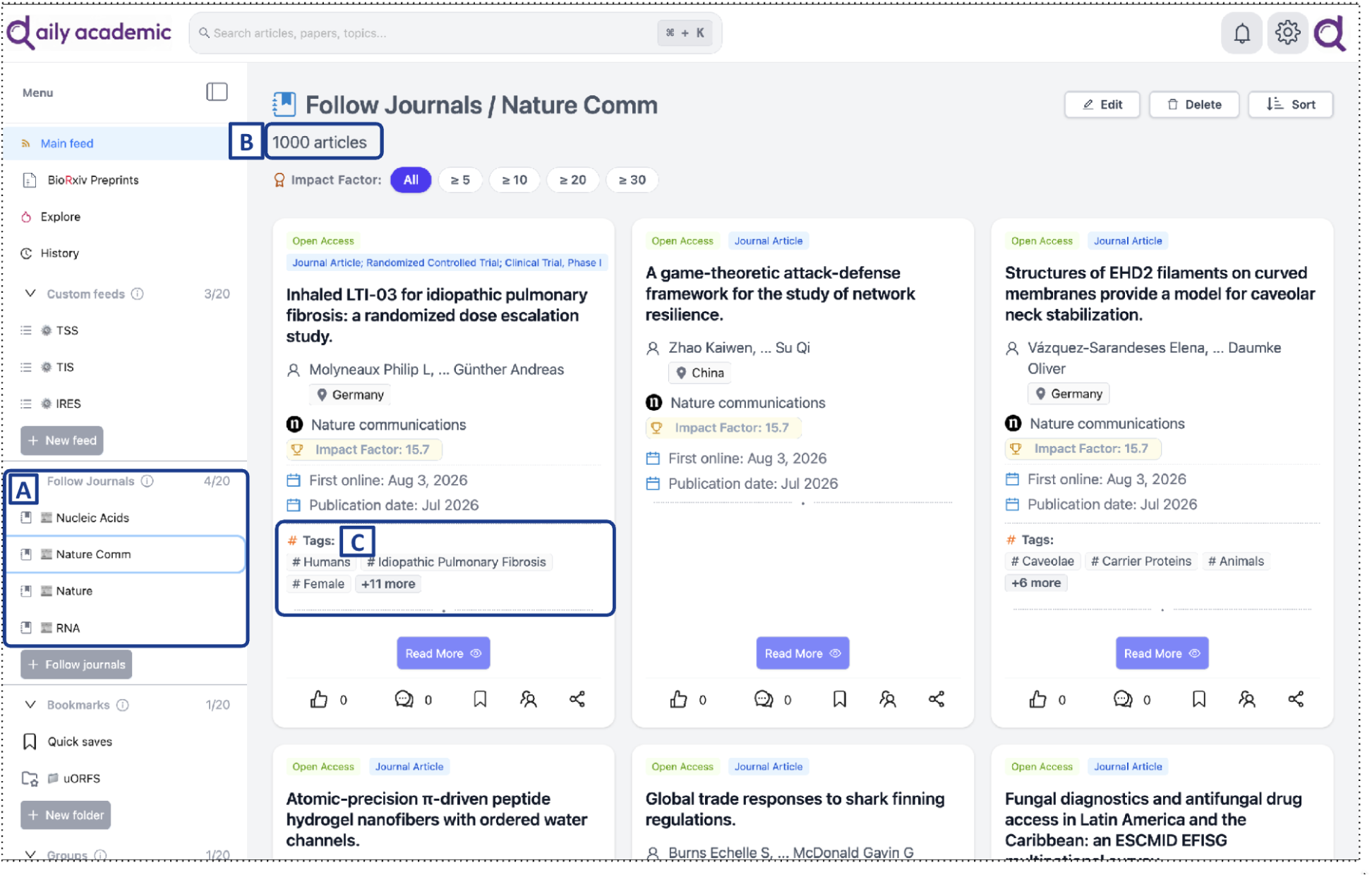
Journal-level following. (A) Followed journals appear in the navigation sidebar. (B) Each journal feed reports the number of indexed articles. (C) Articles are presented with indexed subject terms, supporting rapid assessment of scope. Unlike the main feed, journal feeds are not filtered by the user profile and therefore reflect the full editorial output of the journal.

Tracked journal feeds are integrated into the broader recommendation environment, allowing users to combine source-specific monitoring with personalized semantic discovery within a unified interface. Journal metrics displayed within the platform include impact factor, CiteScore, SJR, h-index, publishing frequency, and acceptance rate. New articles from tracked journals are surfaced alongside personalized recommendations, with smart prioritization applied based on user interest profile and article type preferences.

### Linked Omics Dataset Discovery

Daily-Academic retrieves and displays linked public omics dataset metadata through the Public Omics Explorer (POE) API. Through this API-mediated integration, the platform provides access to structured metadata originating from public omics repositories, including the Gene Expression Omnibus. Each linked dataset entry includes experiment type, organism, disease or phenotype annotations, and accession identifiers such as GSE and SRR records.

Users can navigate bidirectionally between publications and their associated experimental datasets, reducing the need for separate repository searches when investigating specific experimental methodologies, organisms, or disease phenotypes. This integration supports unified retrieval across scientific publications and underlying biological evidence within a single discovery workflow.

### Collaborative and Organizational Features

The platform supports collaborative literature organization through user-defined Groups, enabling researchers to share, annotate, and discuss publications within defined cohorts. Individual article management is supported through saved article collections, topic-based organization, and annotation functionality.

Zotero integration and social sharing functionality are additionally available, supporting export of curated publication sets to external reference management workflows. These features are designed to support both individual literature monitoring and team-based scientific discovery workflows within a unified environment.

### Evaluation Results

#### Offline Retrieval Evaluation Results

Table 4 and Figure 6 summarize overall retrieval performance across the 30 benchmark queries.

**Table 4.** Overall retrieval performance of BM25, PubMedBERT semantic retrieval, and Daily-Academic across 30 biomedical queries. Precision@10 and Precision@30 are reported under the relaxed relevance criterion (labels 1 and 2 treated as relevant); a stricter Precision@10 (label 2 only) is also shown. nDCG@10 and nDCG@30 use graded relevance labels. MeanRel@10 is the average relevance score among the top 10 retrieved articles.

| ***Method*** | ***P@10 (rel≥1)*** | ***P@10 (rel=2)*** | ***nDCG@10*** | ***MeanRel@10*** | ***P@30 (rel≥1)*** | ***nDCG@30*** |
|---|---|---|---|---|---|---|
| *BM25* | *0.723* | *0.260* | *0.565* | *0.983* | *0.656* | *0.773* |
| *PubMedBERT semantic* | *0.580* | *0.117* | *0.613* | *0.697* | *0.466* | *0.733* |
| ***Daily-Academic*** | ***0.913*** | ***0.507*** | ***0.762*** | ***1.420*** | ***0.854*** | ***0.899*** |

Daily-Academic achieved the strongest performance across the benchmark. At rank 10 it reached a Precision@10 of 0.913 when both directly and partially relevant articles were counted, compared with 0.723 for BM25 and 0.580 for the PubMedBERT baseline. Under the stricter criterion (directly relevant only), Daily-Academic obtained 0.507, versus 0.260 for BM25 and 0.117 for PubMedBERT. It also achieved the highest nDCG@10 (0.762 vs 0.565 and 0.613) and the highest mean top-10 relevance, indicating that it surfaced not only more relevant but more directly relevant articles in the highest-ranked positions.

The advantage persisted at a deeper cutoff. At rank 30, Daily-Academic achieved a Precision@30 of 0.854 and an nDCG@30 of 0.899, outperforming BM25 (0.656; 0.773) and PubMedBERT (0.466; 0.733). Among the top 30 results across all queries, Daily-Academic returned the fewest non-relevant and the most directly relevant articles: 422 directly relevant and 131 non-relevant, compared with 192 and 310 for BM25 and 54 and 481 for PubMedBERT (Table 5).

**Table 5.** Distribution of relevance labels among the top-30 retrieved results across all 30 queries. The rel≥1 rate is the proportion of articles with label ≥ 1; the rel=2 rate is the proportion with label = 2.

| ***Method*** | ***Not relevant*** | ***Partially relevant*** | ***Directly relevant*** | ***rel≥1 rate*** | ***rel=2 rate*** |
|---|---|---|---|---|---|
| *BM25* | *310* | *398* | *192* | *0.656* | *0.213* |
| *PubMedBERT semantic* | *481* | *365* | *54* | *0.466* | *0.060* |
| ***Daily-Academic*** | ***131*** | ***347*** | ***422*** | ***0.854*** | ***0.469*** |

Query-level analysis confirmed that these improvements were consistent rather than driven by a few queries. Based on nDCG@10, Daily-Academic ranked first in 19 of the 30 queries (PubMedBERT in 8, BM25 in 3); based on MeanRel@10 it ranked first in 27 of 30, with one BM25 win and two ties (Supplementary Tables S2–S3). Paired Wilcoxon signed-rank tests confirmed the differences were statistically significant across metrics (Table 6); all comparisons remained significant after Bonferroni correction for multiple tests.

**Table 6.** Paired statistical comparison of Daily-Academic against the baseline retrieval methods across the 30 queries (Wilcoxon signed-rank test). Mean differences are the average query-level improvement of Daily-Academic; p-values assess significance.

| ***Comparison*** | ***Metric*** | ***Mean difference*** | ***p-value*** |
|---|---|---|---|
| *Daily-Academic vs BM25* | *nDCG@10* | *+0.197* | *7 × 10<sup>−6</sup>* |
| *Daily-Academic vs PubMedBERT* | *nDCG@10* | *+0.149* | *0.0071* |
| *Daily-Academic vs BM25* | *P@10 (rel≥1)* | *+0.190* | *1.5 × 10<sup>−4</sup>* |
| *Daily-Academic vs PubMedBERT* | *P@10 (rel≥1)* | *+0.333* | *1.7 × 10<sup>−5</sup>* |
| *Daily-Academic vs BM25* | *P@10 (rel=2)* | *+0.247* | *3.7 × 10<sup>−5</sup>* |
| *Daily-Academic vs PubMedBERT* | *P@10 (rel=2)* | *+0.390* | *8 × 10<sup>−6</sup>* |

Overall, the offline evaluation indicates that Daily-Academic produced more effective top-ranked biomedical literature recommendations than both lexical BM25 and PubMedBERT-based semantic retrieval, increasing the proportion of relevant and directly relevant publications among the highest-ranked articles. Because the interaction-driven personalization component is inactive in this user-free setting and the ranking weights were untuned, this advantage is attributable primarily to Daily-Academic’s semantic representation rather than to adaptive personalization, whose value depends on accumulated user interaction and therefore falls outside the scope of this offline benchmark.

## Discussion

The offline benchmark provides evidence that integrating semantic retrieval into a workflow-oriented discovery platform can improve the quality of top-ranked biomedical literature relative to established lexical and dense retrieval baselines. Across 30 queries spanning six biomedical domains, Daily-Academic retrieved a higher proportion of relevant and directly relevant publications at the ranks researchers typically inspect during daily triage, and the advantage was consistent at the query level rather than driven by a small number of favourable topics.

These gains are most meaningful in the context of daily monitoring, where researchers typically inspect only the first few results before moving on. Existing biomedical literature workflows are typically fragmented: researchers maintain separate journal subscriptions, keyword alerts, PubMed saved searches, and reference managers in parallel. Improvements concentrated in the highest ranks therefore reduce the number of irrelevant articles encountered per monitoring session, although the extent to which this reduces researcher workload in practice was not measured in the present study.

### The role of workflow integration

A central design principle of Daily-Academic is the availability of journal tracking, semantic recommendations, and omics dataset discovery within a single interface. This aligns with a broader observation in the information-seeking literature: researchers do not engage in isolated retrieval tasks but in continuous, multi-strategy monitoring workflows that evolve alongside their research interests. Systems designed around individual retrieval tasks require researchers to perform the integration work themselves. Daily-Academic attempts to shift that burden from the user to the platform.

Linked omics dataset discovery is a distinctive capability of the platform, with no direct equivalent in existing literature discovery systems. It is intended to support navigation between published findings and the underlying experimental datasets that informed them - a connection that is otherwise left to the researcher to establish manually across multiple repositories.

### Limitations and caveats

The present evaluation is limited to offline retrieval performance. No user-based evaluation was conducted, so the practical utility of the platform-including its effect on literature monitoring workload, the value of adaptive personalization during sustained use, and the usability of the interface-remains to be established empirically. Statements regarding workflow benefits in this paper therefore reflect design rationale rather than validated outcomes.

Recommendation quality also remains inherently dependent on metadata completeness in the underlying literature sources. Publications with sparse or non-standardized abstracts, or those from journals with limited PubMed indexing coverage, will be less reliably retrieved and ranked. This is a constraint shared by all embedding-based retrieval systems operating over bibliographic metadata rather than full text. Similarly, interdisciplinary topics present challenges when related publications use substantially different terminology across subfields, potentially limiting cross-domain discovery.

The offline benchmark itself has limitations. It comprised 30 queries, which, while spanning diverse monitoring scenarios, may not capture the full variability of user information needs. Relevance judgments were based on titles and abstracts rather than full text, so articles whose decisive evidence appears only in full text may be misjudged. Pooling across all three methods reduces annotation bias but restricts judgments to articles retrieved by at least one evaluated method, so the pool may not contain every relevant article in the corpus. Finally, results reflect a fixed corpus snapshot and may shift as the literature grows. The benchmark also does not evaluate Daily-Academic’s adaptive ranking: with no users the interaction component is inactive and the ranking weights were untuned, so the offline results characterize the platform’s semantic retrieval rather than its personalization. Validating the adaptive ranking framework -for example through a same-encoder ablation with tuned weights, or through interaction-driven evaluation-remains future work. Finally, relevance judgments were produced by a single, non-blinded annotator, which may introduce bias; future evaluations will employ multiple blinded annotators with formal inter-annotator agreement statistics.

The cold-start problem for new users also warrants explicit acknowledgment. The adaptive personalization mechanisms that drive recommendation quality depend on accumulated interaction signals. Newly registered users with no prior activity receive recommendations driven primarily by journal preferences and any uploaded publication history, which may be less precisely tailored than those generated after sustained use. Onboarding mechanisms that help users establish initial interest profiles more efficiently represent an important direction for future development.

### Future directions

User-centered evaluation is a priority for future work. Planned studies include assessment of recommendation relevance and usability with practising biomedical researchers, longer observation periods that allow personalization quality to stabilize, pre/post comparison of monitoring time against baseline workflows, and assessment across research domains beyond bioinformatics. Interaction-driven and simulated-user evaluations are also planned to validate the adaptive ranking framework, which the present user-free benchmark could not assess.

On the platform side, planned developments include richer personalization controls that give users more transparency into why specific publications are recommended, expanded biomedical dataset linkage beyond GEO, and enhanced support for interdisciplinary topic modeling. Further extensions may draw on specialized domain resources and pipelines, including regulatory-element identification tools [27,28,29], translation-initiation and small-ORF analysis frameworks [30] clinical genomic assays [31] and well curated atlases [32,33], reflecting the broader range of biomedical data types that a unified discovery platform could eventually connect. [34]

## Conclusion

Daily-Academic presents a researcher-centered biomedical literature discovery platform that integrates semantic recommendation, structured journal tracking, and linked public omics dataset exploration within a unified scientific monitoring workflow. The platform was designed to reduce fragmentation and operational burden associated with conventional biomedical literature discovery practices while supporting rapid daily scientific content triage.

Offline benchmarking indicates that the platform’s semantic retrieval surfaces a higher proportion of relevant biomedical publications at top ranks than established lexical and dense baselines. By combining semantic retrieval with adaptive personalization and linked dataset exploration, Daily-Academic supports a broader and more context-aware approach to biomedical scientific discovery, although the practical benefits of these workflows remain to be evaluated with users.

As biomedical literature and associated public datasets continue to expand rapidly, workflow-oriented semantic discovery systems may become increasingly important for supporting efficient navigation, monitoring, and interpretation of scientific knowledge across modern biomedical research environments.

## Availability

Daily-Academic is publicly available as a web-based platform at https://www.daily-academic.com. The platform is accessible through a Software-as-a-Service (SaaS) deployment model and is designed to support biomedical literature monitoring and semantic scientific discovery workflows for academic researchers and research organizations. Access to the platform includes researcher-oriented literature discovery, semantic recommendation, journal tracking, and linked public omics dataset exploration functionalities.

## Author contributions

DG conceived the study, designed and implemented the platform, developed the semantic retrieval and recommendation framework, conducted the evaluation, analyzed the data, and wrote the manuscript.

NP contributed scientific supervision to manuscript revision, scientific discussion, conceptual feedback, and critical review of the study and platform design.

All authors reviewed and approved the final manuscript.

## Acknowledgments

The authors thank the researchers and early users who provided feedback during platform development. Dimitris Grigoriadis would like to deeply thank Afroditi Orfanidou for her constant moral encouragement and Kleoniki for her steady inspiration throughout the development of this platform.

## Supplementary Material

**Supplementary Table S1.** The 30 benchmark queries used in the offline retrieval evaluation, adapted from the Public Omics Explorer (POE) evaluation framework [15]. Each original POE query and its Daily-Academic reformulation are shown.

| ID | Category | Original POE query | Daily-Academic query |
|---|---|---|---|
| 1 | Epigenetics | chromatin accessibility in stem cell fate decisions | chromatin accessibility in stem cell fate decisions |
| 2 | Epigenetics | chromatin remodeling in cancer initiation | chromatin remodeling mechanisms in cancer initiation |
| 3 | Epigenetics | epigenetic control of lineage commitment | epigenetic regulation of lineage commitment |
| 4 | Epigenetics | epigenetic resetting in reprogramming | epigenetic resetting during cellular reprogramming |
| 5 | Immune regulation & cancer | inflammatory signaling in cardiovascular disease | inflammatory signaling mechanisms in cardiovascular disease |
| 6 | Immune regulation & cancer | metabolic reprogramming in immune activation | metabolic reprogramming during immune cell activation |
| 7 | Immune regulation & cancer | nutrient sensing pathways in transcriptional regulation | nutrient-sensing pathways in transcriptional regulation |
| 8 | Immune regulation & cancer | regulatory T cell suppressive mechanisms | regulatory T cell suppressive mechanisms in immune regulation |
| 9 | Immune regulation & cancer | stress response pathways in cancer survival | stress response pathways promoting cancer cell survival |
| 10 | Neurological disease | gene expression changes in Parkinson’s disease | gene expression changes in Parkinson’s disease |
| 11 | Neurological disease | neuroinflammation signatures in Alzheimer’s disease | neuroinflammation signatures in Alzheimer’s disease |
| 12 | Neurological disease | transcriptional dysregulation in Parkinson’s disease | transcriptional dysregulation in Parkinson’s disease |
| 13 | Non-coding RNA biology | microRNA dysregulation in cancer | microRNA dysregulation in cancer |
| 14 | Non-coding RNA biology | miRNA promoter variants in cancer | miRNA promoter variants in cancer-associated gene regulation |
| 15 | Non-coding RNA biology | mutations in miRNA promoter regions | mutations in miRNA promoter regions and regulatory effects |
| 16 | Non-coding RNA biology | non-coding RNA influence on transcriptional regulation | non-coding RNA effects on transcriptional regulation |
| 17 | Non-coding RNA biology | RNA editing in neuronal function | RNA editing in neuronal function |
| 18 | Non-coding RNA biology | transcription factors binding to miRNA promoters | transcription factor binding at miRNA promoters |
| 19 | Non-coding RNA biology | transcriptional regulation of miRNA promoter regions | transcriptional regulation of miRNA promoter regions |
| 20 | Regulatory genomics | cross-talk between transcription factors and signaling pathways | cross-talk between transcription factors and signaling pathways |
| 21 | Regulatory genomics | long-range chromatin contacts in transcription control | long-range chromatin interactions in transcriptional control |
| 22 | Regulatory genomics | long-range chromatin interactions in hematopoietic cells | long-range chromatin interactions in hematopoietic cells |
| 23 | Regulatory genomics | mechanisms of transcriptional memory | molecular mechanisms of transcriptional memory |
| 24 | Regulatory genomics | ribosome profiling of translation initiation events | ribosome profiling of translation initiation events |
| 25 | Regulatory genomics | stochastic gene expression in single cells | stochastic gene expression in single cells |
| 26 | Regulatory genomics | transcription factor binding dynamics | transcription factor binding dynamics in gene regulation |
| 27 | Regulatory genomics | transcription factor mutation effects on target networks | transcription factor mutations and target gene networks |
| 28 | Stem cells and development | adaptive gene expression in response to environment | adaptive gene expression responses to environmental signals |
| 29 | Stem cells and development | gene regulatory networks in stem cells | gene regulatory networks in stem cells |
| 30 | Stem cells and development | pluripotency transcriptional networks | transcriptional networks controlling pluripotency |

**Supplementary Table S2.**
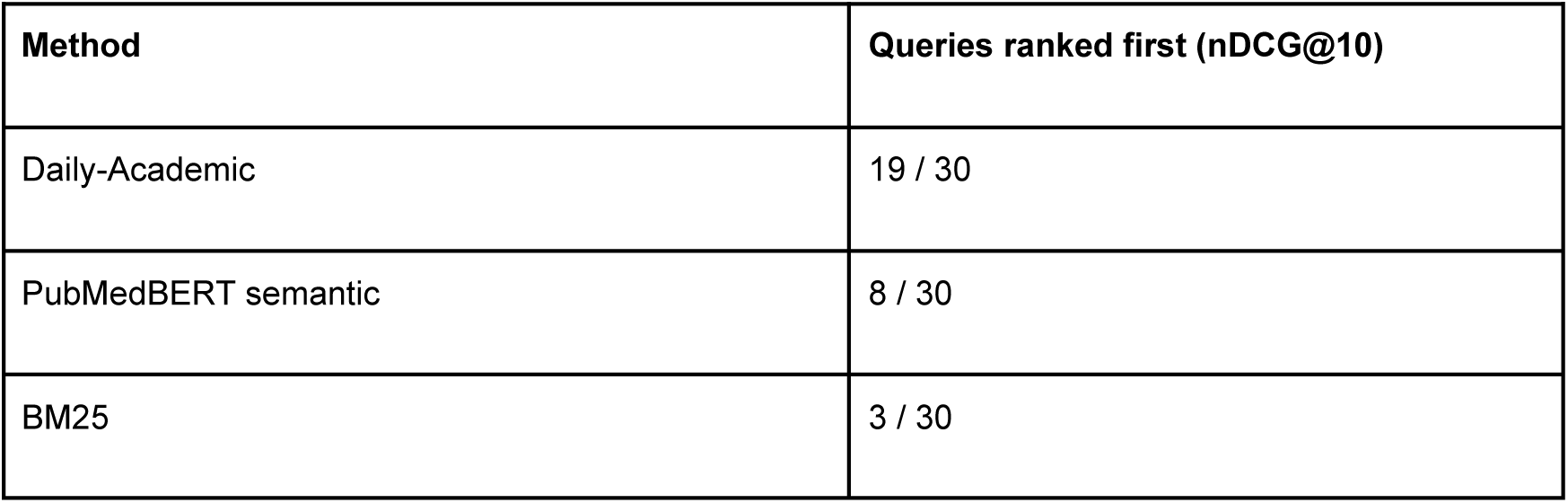
Query-level comparison based on nDCG@10. For each query, the method with the highest nDCG@10 score was identified.

| **Method** | **Queries ranked first (nDCG@10)** |
|---|---|
| Daily-Academic | 19 / 30 |
| PubMedBERT semantic | 8 / 30 |
| BM25 | 3 / 30 |

**Supplementary Table S3.** Query-level comparison based on MeanRel@10. For each query, the method with the highest mean relevance score among the top 10 retrieved articles was identified; ties indicate equal top scores.

| **Method** | **Queries ranked first (MeanRel@10)** |
|---|---|
| Daily-Academic | 27 / 30 |
| BM25 | 1 / 30 |
| PubMedBERT semantic | 0 / 30 |
| Ties (same score) | 2 / 30 |

---

**Supplementary Figure S1.**
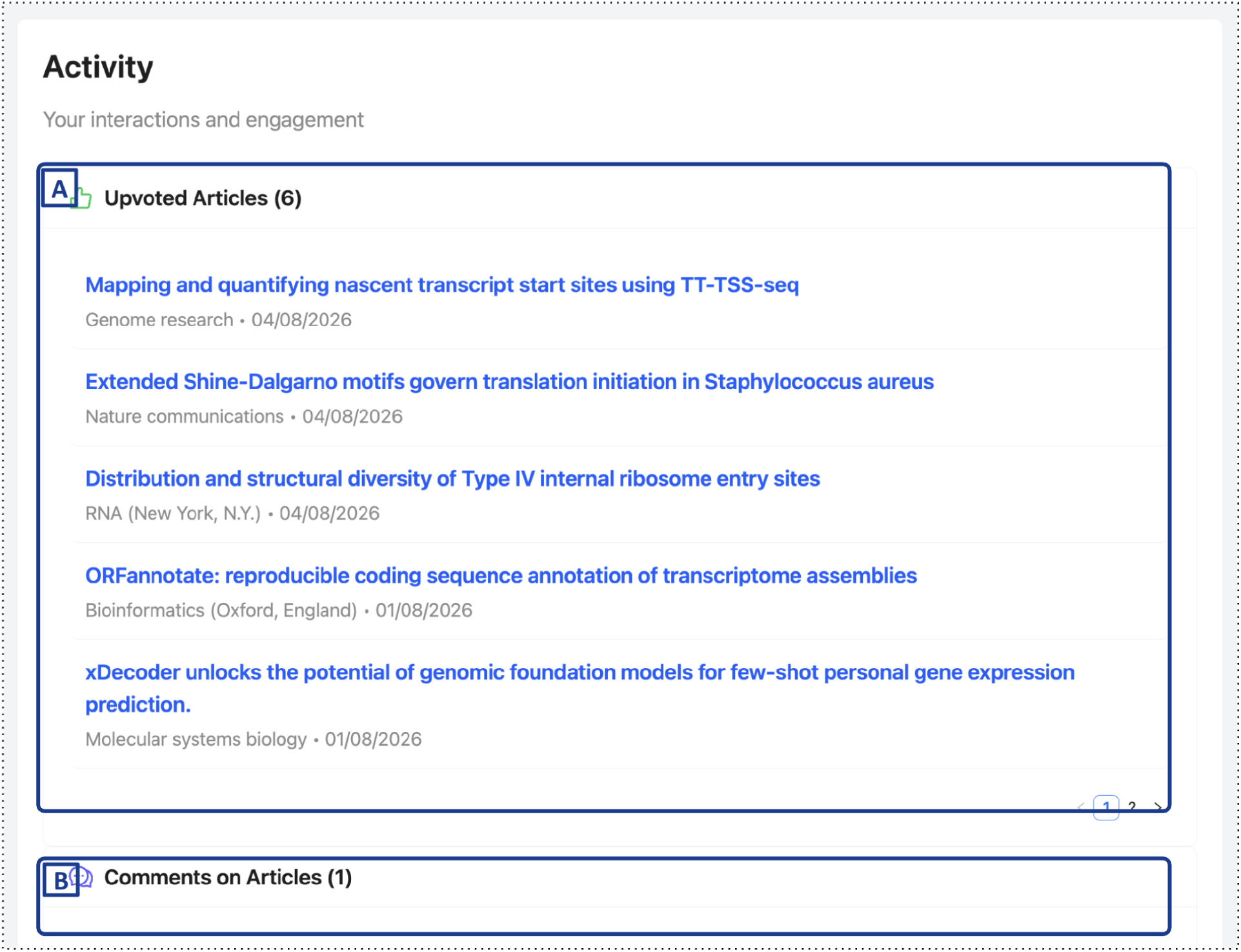
Interaction history view. (A) Articles the user has upvoted, listed with source journal and date. (B) Comments the user has posted. When the adaptive personalization option is enabled (Fig. 3A), upvotes are incorporated into the daily update of the user profile vector, providing an implicit personalization signal alongside the explicitly stated research interests.

